# Altered Baseline Brain Network Topology in High-Risk Individuals Progressing to Mild Cognitive Impairment

**DOI:** 10.64898/2026.02.05.704129

**Authors:** Hema Nawani, Ranjith Jaganathan, Veeky Baths, The PREVENT-AD Research Group

## Abstract

**Background:** Identifying early brain-based markers of cognitive decline is critical for preventive strategies in Alzheimer’s disease. Individuals with a familial risk may exhibit subtle functional brain changes years before clinical symptoms emerge. This exploratory study examined whether baseline functional brain network topology differentiates high-risk cognitively normal older adults who later progress to mild cognitive impairment (MCI) from those who remain cognitively stable.

**Methods:** Baseline resting-state fMRI data were analyzed from 90 cognitively normal adults with a family history of Alzheimer’s (PREVENT-AD cohort), classified longitudinally as converters (MCI-C; n=45) or non-converters (MCI-NC; n=45). Whole-brain functional networks were analyzed across multiple thresholds; primary results are reported at 12% network density, with robustness verified at 16% density. Group differences were assessed using ANCOVA or Rank ANCOVA (controlling for age, sex, and education) at an uncorrected threshold (p < 0.05). Predictive utility was evaluated via a 100-repetition nested cross-validation machine-learning framework on a multimodal feature set combining functional network metrics, average cortical thickness, and plasma p-tau217, with covariates included within training folds.

**Results:** At baseline, MCI-C participants were older, had fewer years of education, exhibited higher plasma p-tau217 levels, and showed trend-level lower MoCA scores. At 12% density, MCI-C showed increased average nodal strength (F=4.50, p=0.036, ηp^2^=0.050) and reduced global efficiency (F=4.07, p=0.046, ηp^2^=0.045). Increased betweenness centrality within the Default Mode Network (F=4.07, p=0.046, ηp^2^=0.045) and trend-level increases in average clustering (F=3.10, p=0.081, ηp^2^=0.035) were observed. Initial largest connected component (LCC) showed a trend-level decrease (F=3.84, p=0.053, ηp^2^=0.043).

At 16% density, MCI-C exhibited significantly reduced initial LCC (F=4.41, p=0.038, ηp^2^=0.049) and increased nodal strength (F=4.29, p=0.041, ηp^2^=0.048), with directionally consistent trend-level reductions in global efficiency (F=3.74, p=0.056, ηp^2^=0.042). In machine learning, the k-nearest neighbors classifier showed the most stable performance (nested CV accuracy=59.6%; test F1-score=0.56). Feature stability analysis identified global efficiency (selected in 25.8% of iterations) and critical drop (19.4%) as the most consistent predictors.

**Conclusion:** Baseline disruptions in functional network integration precede clinical conversion to MCI. The consistent selection of graph-theoretical metrics, particularly global efficiency and critical drop, as top predictors suggests that functional network reorganization provides unique information for classification before widespread cortical atrophy emerges.

## 1. Introduction

Mild Cognitive Impairment (MCI) represents a transitional stage between healthy aging and dementia, most commonly Alzheimer’s disease (AD). It is characterized by measurable but not yet functionally disabling cognitive decline and carries a substantially increased risk of conversion to dementia^1,2^. The risk of progression is further elevated by advanced age, vascular comorbidity, genetic predisposition (e.g., APOE ε4 allele), and a family history of neurodegenerative disease^3,4^. A major goal in preventative neurology is the identification of early neurobiological markers in high-risk individuals before clinical conversion^5^. Initiatives such as the Pre-symptomatic Evaluation of Experimental or Novel Treatments for AD (PREVENT-AD) program provide an opportunity to investigate brain changes in high-risk individuals (cognitively unimpaired older individuals with a parental or multiple-sibling history of AD) before clinical conversion^6^.

Resting-state functional MRI (rs-fMRI) offers a non-invasive approach to characterize the brain’s intrinsic functional organization. Spontaneous low-frequency fluctuations in the blood-oxygen-level–dependent (BOLD) signal reveal large-scale functional networks^7^. Graph theory provides a formal mathematical framework to describe their topological properties, including efficiency, clustering, modularity, and hub structure^8^. Prior studies have demonstrated that MCI and AD are associated with disrupted network topology compared to cognitively normal controls, often reflected in reduced efficiency, increased path length, or impaired hub connectivity^9,10^.

Beyond traditional static measures, network resilience based modeling provides a dynamic perspective on brain vulnerability^11^. By simulating targeted attack on highly connected nodes, these approaches estimate extent of network disintegration and at what point catastrophic failure occurs^11,12,13^. Following the resilience procedure introduced by Argiris et al. (2024)^11^, we simulated targeted network attack by iteratively removing nodes ranked by descending nodal strength and computing the size of the largest connected component (LCC) after each removal. Brain resilience was quantified as the critical drop, defined as the minimum slope (largest single-step decrease) in the LCC decay curve during targeted attack. Given that AD preferentially affects high-centrality and high-metabolic-demand hub regions, particularly within the default mode network, this model provides a mechanistically grounded proxy for early network vulnerability^11^. This framework may be particularly relevant in AD, where hub regions within networks such as the default mode network are known to accumulate early pathology^14^.

In this exploratory study, we analyzed baseline resting-state fMRI from high-risk older adults in the PREVENT-AD cohort and compared individuals who subsequently progressed to mild cognitive impairment (MCI; “converters”: MCI-C) with those who remained cognitively stable over a five-year clinical follow-up (“non-converters”: MCI-NC). We applied graph-theoretical and resilience analysis to characterize alterations in functional brain network topology during the pre-symptomatic phase. Based on prior work in Alzheimer’s disease, we hypothesized that MCI-C would exhibit reduced global efficiency, altered hub-level centrality, and lower network resilience, quantified by a more negative critical drop (largest single-step decrease) in the largest connected component (LCC) during simulated targeted attack reflecting increased functional network breakdown compared with MCI-NC.

## 2. Methods

### 2.2 Participants and Study Design

Data used in preparation of this article were obtained from the PRe-symptomatic EValuation of Experimental or Novel Treatments for Alzheimer’s Disease (PREVENT-AD) registered repository, available at https://registeredpreventad.loris.ca. The Investigators of the PREVENT-AD program contributed to the PREVENT-AD provided data but did not participate in data analysis or writing of this report. This study analyzed baseline resting-state functional MRI (rs-fMRI) and T1-weighted structural MRI data from the PREVENT-AD cohort. Participants were identified as high-risk based on a confirmed family history of Alzheimer’s disease. Clinical follow-up data were used to categorize participants into two groups:

Converters (MCI-C, n = 45): Participants who progressed to mild cognitive impairment (MCI) during follow-up. Non-converters (MCI-NC, n = 45): participants who remained cognitively stable throughout follow-up. Only baseline neuroimaging data collected prior to any clinical conversion were analyzed.

### 2.2 MRI Preprocessing

Preprocessing was performed with fMRIPrep (version 24.1.1), integrating FSL, FreeSurfer, ANTs, and AFNI^16^. **Structural images:** Intensity non-uniformity correction (N4BiasField), skull stripping, tissue segmentation (gray matter, white matter, cerebrospinal fluid), cortical surface reconstruction (FreeSurfer), and nonlinear registration to MNI152NLin2009cAsym space. **Functional images:** The first four volumes were discarded to allow for signal stabilization. Preprocessing included slice-timing correction, motion correction, susceptibility distortion correction, and alignment to the native T1-weighted space using boundary-based registration. Functional data were normalized to the MNI152NLin2009cAsym template (MNI space) and resampled to 2mm isotropic resolution. **Denoising and filtering:** Physiological noise components were removed using aCompCor from WM and CSF compartments. The time series were detrended, z-standardized, and band-pass filtered (0.01–0.09 Hz). **Parcellation and Functional Network Construction:** BOLD time series were extracted from 421 brain regions, consisting of: Cortical parcellation: Schaefer 400-region, 17-network parcellation^17^ and Subcortical regions: Harvard– Oxford probabilistic atlas thresholded at 50%^18^. Functional connectivity (FC) matrices were generated as Pearson correlation coefficients between all regions. Negative correlations were set to zero, and the matrices were Fisher z-transformed. To maintain consistent network density, the top 12% of positive connections were retained across all subjects, producing weighted, undirected graphs constructed with NetworkX^19^. Graph-Theoretical Metrics Global and nodal metrics were computed and categorized as shown in Figure 1. Network resilience was quantified following Argiris et al.^11^ using a targeted attack framework in which nodes were iteratively removed in descending order of nodal strength, and the size of the largest connected component (LCC) was tracked after each removal. The critical drop or resilience was operationally defined as the iteration of the steepest slope in the largest connected component (LCC) decay.

**Figure 1.**
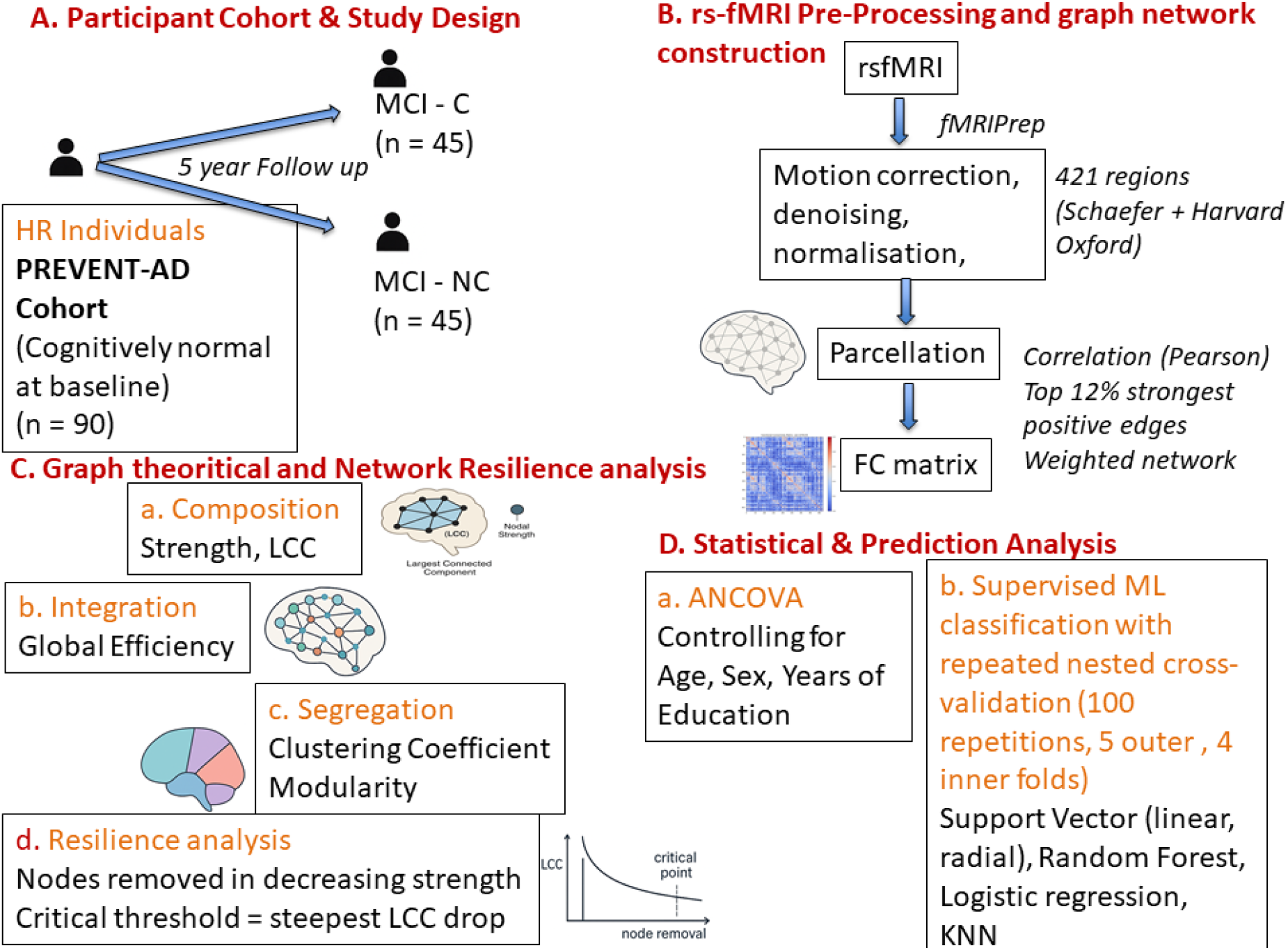
Overview of study design, preprocessing pipeline, graph-theoretical metrics, and statistical analysis. **(A)** Participant cohort and study design. Ninety cognitively normal, high-risk individuals from the PREVENT-AD cohort were followed for approximately 5 years and subsequently classified as converters to Mild Cognitive Impairment (MCI-C ; n=45) or non-converters (MCI-NC ; *n*=45). **(B)** Resting-state fMRI preprocessing pipeline implemented with fMRIPrep, including motion correction, spatial normalization, and denoising. Time series were extracted from 421 brain regions (400 cortical Schaefer parcels and 21 Harvard-Oxford subcortical regions) to generate individual functional connectivity (FC) matrices. **(C)** Graph-theoretical and network resilience analysis framework. Composition metrics included nodal strength and the size of the largest connected component (LCC). Network integration was assessed using global efficiency, while segregation was evaluated via clustering coefficient and modularity. Resilience was estimated by simulating targeted removal of high-strength nodes and quantifying LCC decay to derive the critical drop. **(D)** Statistical and predictive modeling. Group differences were evaluated using ANCOVA controlling for age, sex, and years of education. Supervised machine learning models (support vector classification, random forest, logistic regression, and k-nearest neighbors) were trained to predict conversion from baseline network metrics.

### 2.3 Statistical Analysis

Demographic and clinical variables, including age, sex, years of education, and Montreal Cognitive Assessment (MoCA) scores, were obtained are summarized in Table 2.

**Table 1.** Graph-theoretical measures categorized by functional domain. *Composition* describes basic structural features of the network. *Centrality* reflects nodal influence. *Integration* and *segregation* characterize global and local information flow, respectively. *Resilience* metrics assess robustness under simulated targeted attacks

| **Composition** | **Integration** | **Centrality** | **Segregation** | **Resilience** |
| --- | --- | --- | --- | --- |
| Strength | Global efficiency | Betweenness centrality* | Clustering coefficient | Critical drop |
| Initial LCC size |  | Eigenvector centrality* |  |  |

\* computed within Default Mode Network (DMN) networks parcels

**Table 2.** Group Comparisons between MCI-NC and MCI-C on demographic, cognitive, and biomarker variables.

| Variable | MCI Non-Converter, Mean (SD) | MCI Converter, Mean (SD) | Mann–Whitney U | Z | *p*-value | Effect Size (*r*) |
| --- | --- | --- | --- | --- | --- | --- |
| Age (months) | 751.64 (61.74) | 789.71 (73.59) | 1354.5 | 2.76 | 0.006 | 0.29 |
| Education (years) | 15.62 (2.57) | 14.20 (3.26) | 705 | -2.5 | 0.012 | 0.26 |
| MOCA total score | 28.09 (1.69) | 27.56 (1.62) | 793.5 | -1.8 | 0.071 | 0.19 |
| ptau217 | 2.29 (1.56) | 2.57 (1.05) | 1315 | 2.44 | 0.015 | 0.26 |
| Mean FD | 0.25 (0.11) | 0.27 (0.12) | 1095 | 0.67 | 0.506 | 0.07 |

Between-group differences (MCI-C vs. MCI-NC) were tested using ANCOVA, controlling for age, sex, and years of education. Results were reported as uncorrected p-values (p < 0.05) consistent with the exploratory design, with partial η^2^ provided as effect sizes. Benjamini–Hochberg false discovery rate (FDR) correction was applied post hoc across graph-theoretical metrics. To mitigate concerns regarding the arbitrariness of thresholding, all graph-theoretical analysis were repeated at two network densities (12%, and 16%), spanning a commonly used range in resting-state functional connectivity studies.

### 2.4 Machine Learning Analysis

A supervised machine learning framework was implemented in MATLAB R2024b to evaluate the predictive utility of baseline multimodal features for conversion from high risk to mild cognitive impairment (MCI). The feature set comprised functional brain network metrics, including initial largest connected component (LCC), critical drop during targeted attack, global efficiency, average clustering coefficient, modularity, and average nodal strength, along with a structural measure (mean cortical thickness) and the plasma biomarker p-tau217. To ensure unbiased performance estimation and minimize overfitting, a repeated nested cross-validation (CV) framework was employed, consisting of 100 repetitions with 5 outer folds and 4 inner folds, using stratified partitioning to preserve class balance. Prior to nested CV, the dataset was split into a fixed training set (80%) and an independent hold-out test set (20%). Within each outer training fold, the effects of age, sex, and years of education were regressed out from the feature matrix using linear regression, and the resulting regression coefficients were applied to the corresponding validation data to prevent information leakage. Features were ranked within each outer training fold only using a two-sample t-test to define an ordered feature list for dimensionality reduction; importantly, this ranking was used solely for feature ordering and did not constitute hypothesis testing or group-level statistical inference. The inner CV loop was used to jointly optimize model hyperparameters and the number of top-ranked features, with selection based on the highest average F1 score across inner validation folds. Using the optimal configuration, models were trained on each outer training fold and evaluated on the corresponding outer validation fold. Performance metrics were aggregated across all repetitions, yielding the mean and standard deviation of outer CV accuracy, as well as the most frequently selected hyperparameters and feature counts for each model. Final models were subsequently trained on the full training dataset using the most frequently selected hyperparameters and feature counts derived from nested CV. Covariates were regressed out, top-ranked features were selected using training data only, and model performance was evaluated on the independent 20% test set, reporting accuracy and F1 score.

## 3. Results

### 3.1 Demographic and Clinical Characteristics

Baseline demographic, clinical, and biomarker data are summarized in Table 2. The group of individuals who later converted to MCI (MCI-C) was significantly older than the stable non-converter group (MCI-NC) (U = 1354.5, p = 0.006, r = 0.29) and had fewer years of formal education (U = 705, p = 0.012, r = 0.26). Plasma phosphorylated tau-217 (ptau217) levels were also significantly elevated in the MCI-C group (U = 1315, p = 0.015, r = 0.26). No significant differences were found in mean framewise displacement (U = 1095, p = 0.506). A trend-level difference was observed in baseline Montreal Cognitive Assessment (MoCA) scores (U = 793.5, p = 0.071), with converters showing lower scores (MCI-C: 27.56 ± 1.62; MCI-NC: 28.09 ± 1.69).

### 3.2 Graph-Theoretical Network Characteristics

Group comparisons of graph-theoretical metrics were conducted using ANCOVA or Rank ANCOVA, controlling for age, sex, and education. Robustness analysis conducted at two network densities (12%, and 16%). The results from 0.12 density are considered in Table 3 and results from 0.16 density are presented in supplementary material. To account for multiple comparisons across graph-theoretical metrics, Benjamini–Hochberg false discovery rate (FDR) correction was applied. As no effects survived FDR correction, uncorrected p-values are reported to characterize effect sizes and directional trends in this exploratory analysis.As detailed in Figure 2 and Table 3, this revealed specific topological differences between converters and non-converters at baseline. The MCI-C group demonstrated significantly reduced global efficiency compared to MCI-NC (F = 4.08, p =.0466, η^2^ = 0.046). Concurrently, converters exhibited significantly increased average nodal strength (F = 4.50, p =.0367, η^2^ = 0.050). Betweeness DMN was increased in MCI-C compared to MCI-NC (F = 4.07, p =.0466, η^2^ = 0.045)

**Table 3.**
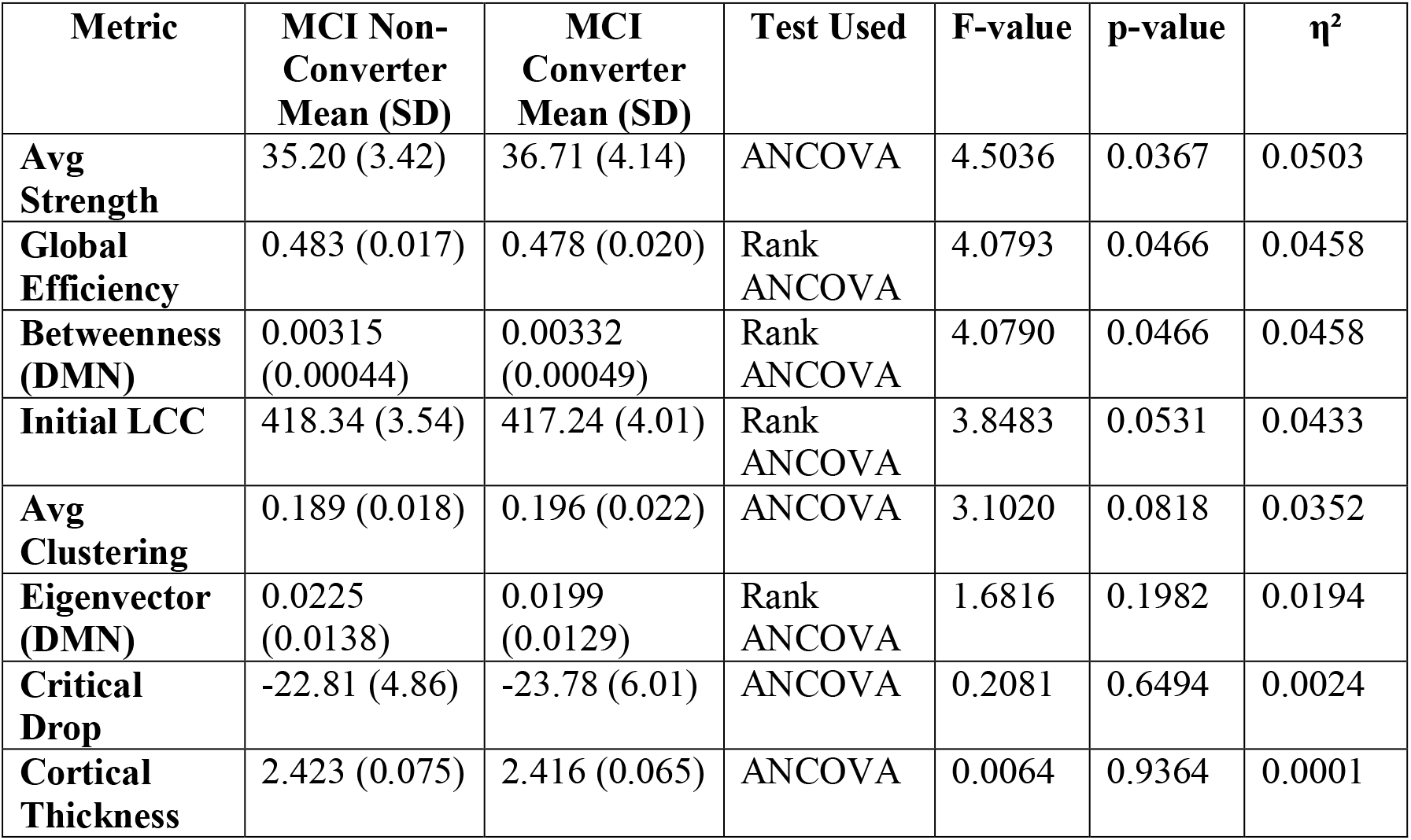
Group differences in graph-theoretical metrics at network density of 12% in baseline high-risk individuals who later converted to MCI (MCI-C) and those who remained cognitively stable (MCI-NC). Uncorrected p-values are shown. After Benjamini–Hochberg false discovery rate (FDR) correction across graph-theoretical metrics, none of the effects remained significant.

| **Metric** | **MCI Non-Converter Mean (SD)** | **MCI Converter Mean (SD)** | **Test Used** | **F-value** | **p-value** | **$\eta^2$** |
| --- | --- | --- | --- | --- | --- | --- |
| **Avg Strength** | 35.20 (3.42) | 36.71 (4.14) | ANCOVA | 4.5036 | 0.0367 | 0.0503 |
| **Global Efficiency** | 0.483 (0.017) | 0.478 (0.020) | Rank ANCOVA | 4.0793 | 0.0466 | 0.0458 |
| **Betweenness (DMN)** | 0.00315 (0.00044) | 0.00332 (0.00049) | Rank ANCOVA | 4.0790 | 0.0466 | 0.0458 |
| **Initial LCC** | 418.34 (3.54) | 417.24 (4.01) | Rank ANCOVA | 3.8483 | 0.0531 | 0.0433 |
| **Avg Clustering** | 0.189 (0.018) | 0.196 (0.022) | ANCOVA | 3.1020 | 0.0818 | 0.0352 |
| **Eigenvector (DMN)** | 0.0225 (0.0138) | 0.0199 (0.0129) | Rank ANCOVA | 1.6816 | 0.1982 | 0.0194 |
| **Critical Drop** | -22.81 (4.86) | -23.78 (6.01) | ANCOVA | 0.2081 | 0.6494 | 0.0024 |
| **Cortical Thickness** | 2.423 (0.075) | 2.416 (0.065) | ANCOVA | 0.0064 | 0.9364 | 0.0001 |

**Figure 2.**
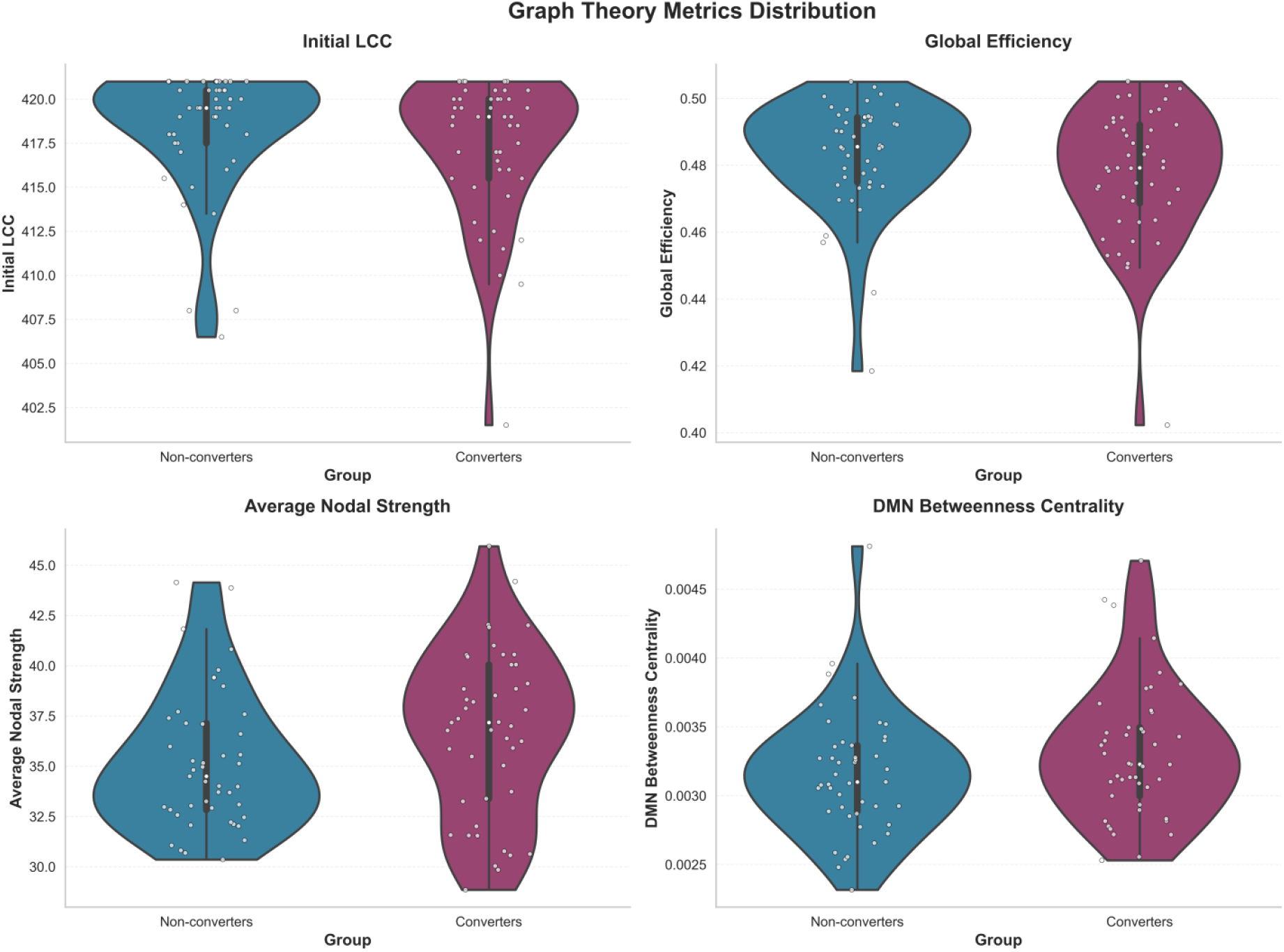
Distribution of key graph-theoretical metrics in high-risk individuals who converted (Converters) and those who remained cognitively stable (Non-converters).

**Figure 3.**
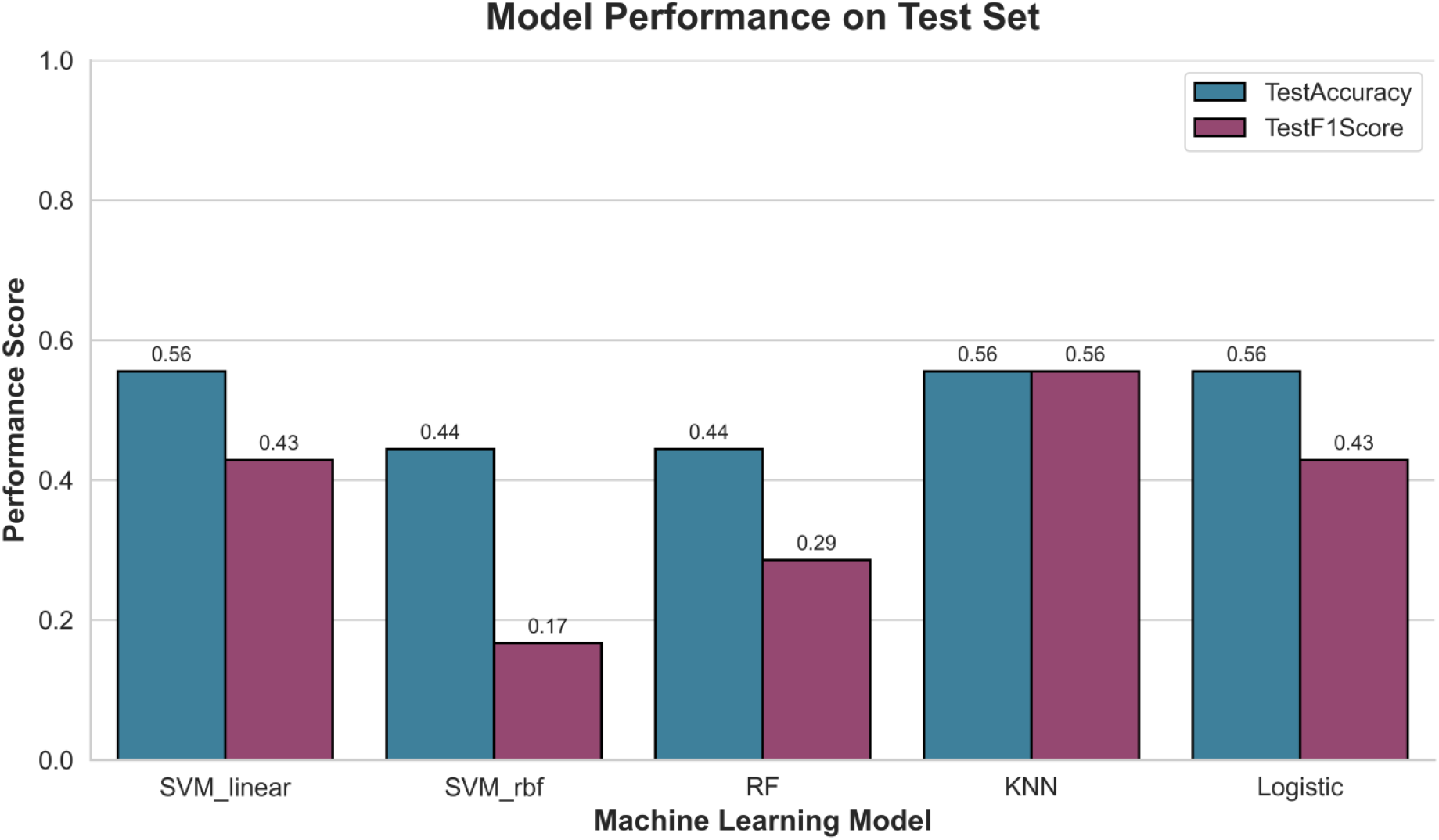
Model Performance on Test Set

Trend-level increases were observed in MCI-C group compared to MCI-NC for the initial largest connected component (LCC) size (F = 3.85, p =.0531, η^2^ = 0.043) and average clustering coefficient (F = 3.10, p =.0818, η^2^ = 0.035). No other graph-theoretical metrics showed statistically significant differences between groups after covariate adjustment (p <.05 uncorrected for multiple comparisons).

This figure presents the predictive performance of various supervised machine learning models (KNN, Logistic Regression, SVM, and Random Forest) used to identify individuals who would later convert to Mild Cognitive Impairment (MCI) based on their baseline brain network metrics, average cortical thickness, and plasma p-tau217.

### 3.3 Network Resilience

Targeted attack simulations revealed no significant difference in network resilience between groups, as measured by the critical drop (F = 0.20, p =.6494, η^2^ = 0.002). Importantly, no group differences emerged for network resilience metrics at any density. Effect sizes and directions were stable across thresholds, indicating that the observed pattern of inefficient hyperconnectivity and preserved resilience is not driven by density selection (Supplementary Tables S1).

### 3.4 Predictive Modeling of MCI Conversion

The supervised machine learning framework was implemented to evaluate the predictive utility of baseline network metrics for MCI conversion. The best performance, on the independent hold-out test set, was achieved by the k-Nearest Neighbors classifier, which attained an accuracy and F1-score of 0.56 and mean validation accuracy of 0.59 in outer CV. Feature stability analysis identified global efficiency (selected in 25.8% of iterations) and critical drop (19.4%) as the most consistent predictors.

## 4. Discussion

### 4.1 Baseline Characteristics Differentiate Future Converters from Stable Individuals

At baseline, clear distinctions were observed between high-risk individuals who would later progress to mild cognitive impairment (MCI-C) and those who remained cognitively stable (MCI-NC). The MCI-C group was significantly older and had fewer years of formal education than the MCI-NC group, two established demographic risk factors for cognitive decline. Biologically, the MCI-C group exhibited significantly higher plasma concentrations of p-tau217, a core biomarker of Alzheimer’s pathology strongly linked to clinical progression^20,21^. A trend-level lower MoCA score further suggested that subtle cognitive inefficiencies were already emerging in the MCI-C group, despite both groups being classified as cognitively normal. Critically, the groups were comparable in terms of sex distribution and head motion (mean framewise displacement), strengthening the interpretation that the observed functional network differences are related to underlying pathology rather than demographic imbalances or motion artifacts.

In summary, the MCI-C group presented with a profile defined by older age, less education, higher p-tau217, and a trend toward lower cognitive performance that differentiated them at baseline from their stable counterparts and is consistent with a greater underlying biological vulnerability to progression.

### 4.2 Increased Strength, Reduced Global Efficiency, and Elevated Local Clustering: The Phenotype of Inefficient Hyperconnectivity

A central finding of the present study is the co-occurrence of increased average nodal strength, reduced global efficiency, and a trend-level increase in local clustering coefficient in individuals who later progressed to MCI-C. Together, this triad defines a phenotype of *inefficient hyperconnectivity*, in which local functional coupling is strengthened while the capacity for fast, integrative communication across the whole brain is compromised.

From a graph-theoretical perspective, these findings are internally consistent. Average nodal strength reflects the overall magnitude of functional coupling and is sensitive to increases in local synchrony. Clustering coefficient indexes the tendency of neighboring nodes to form tightly interconnected local neighborhoods, whereas global efficiency captures the inverse of characteristic path length and reflects the efficiency of long-range information transfer. Thus, increased nodal strength and clustering can coexist with reduced global efficiency when enhanced local connectivity is accompanied by disrupted or inefficient long-range integration. In this context, an increase in clustering does not imply preserved network efficiency; rather, it may signal a shift toward more locally segregated processing at the expense of global communication.

This dissociation aligns with theoretical models of Alzheimer’s disease (AD) network reorganization that propose an early phase of hyperconnectivity followed by progressive network failure^22^. Importantly, Jones et al^22^ present a systems-level framework of cascading hub overload, illustrating how highly connected hub regions may initially increase their functional engagement but eventually become vulnerable to overload and breakdown. As this work does not empirically report whole-brain graph metrics such as nodal strength, clustering coefficient, or global efficiency, it is cited here strictly as a conceptual precedent rather than a metric-specific comparison.

#### 4.2.1 Preclinical Compensation versus Pathological Disruption

In preclinical and at-risk cohorts, network alterations are often interpreted as compensatory rather than degenerative. For example, Mormino et al.^23^ demonstrated that cognitively normal older adults with higher amyloid burden show region-specific increases in default mode network (DMN) functional connectivity, particularly in dorsal and anterior medial prefrontal and lateral temporal cortices with decreases in connectivity within episodic memory related DMN hubs. This pattern was interpreted as early compensatory reorganization rather than uniform hyperconnectivity. Similarly, Arenaza-Urquijo et al.^24^ reported increased functional connectivity between the anterior cingulate cortex and distributed frontal, temporal, and parietal regions in cognitively normal elders with higher educational attainment, consistent with reserve related reinforcement of long-range coupling in the absence of overt pathology.

In our familial-risk cohort, individuals who did not convert to MCI exhibited significantly higher educational attainment than converters, suggesting that cognitive reserve may modulate the balance between compensatory coupling and pathological network breakdown. In this framework, increased functional coupling may initially support performance, but its effectiveness depends on whether global integration remains intact.

##### Graph-Theoretical Evidence across MCI and AD

In contrast to preclinical compensation, graph-theoretical studies in symptomatic MCI and AD more consistently indicate network disintegration. Seminal resting-state fMRI work by Supekar et al.^25^ reported reduced clustering coefficient in AD, without a concomitant change in average path length, suggesting early disruption of local network organization. Conversely, Sanz-Arigita et al.^26^ observed decreased characteristic path length with preserved clustering in AD, indicating that metric-specific findings vary substantially across studies, likely due to differences in preprocessing, thresholding, and cohort characteristics.

Across more recent rs-fMRI studies in MCI, reductions in global efficiency emerge as one of the most robust findings, whereas changes in the clustering coefficient remain heterogeneous. Crucially, a recent analysis comparing stable and progressive MCI found that significant reductions in the small-world parameter and clustering coefficient were primarily observed in the progressive MCI group, while the stable group did not show significant global property declines^27^. This finding emphasizes that the loss of efficient local and global topology is strongly linked to conversion. Large-scale progression studies reinforce this dissociation between integration and segregation. This finding underscores that disruption of both global integration and local specialization is closely linked to clinical conversion. In aMCI, increased characteristic path length and reduced overall connectivity reflect impaired large-scale integration and inefficient information transfer across distributed brain systems, consistent with canonical models of functional brain organization^28, 29^. Xiang et al.^30^ demonstrated increased characteristic path length in MCI, reflecting reduced network integration, whereas Brier et al.^31^ reported progressive loss of clustering and network segregation across the continuum from cognitively normal individuals to MCI and AD, without a significant change in path length. Together, these findings indicate that disruption of integrative capacity whether indexed by increased path length, reduced global efficiency, or loss of network segregation is a consistent marker of early network breakdown, even when individual graph metrics show variable behavior. This interpretation is further supported by our biomarker findings. Elevated baseline p-tau217 a marker closely linked to tau deposition and propagation provides a plausible biological mechanism for the observed dissociation between local and global network properties. Tau pathology has been shown to preferentially disrupt long-range connectivity and hub integrity^32, 33^, potentially forcing the network toward increased local coupling while undermining integrative efficiency.

### 4.3 Increased DMN Betweenness Centrality Suggests Hub Overload

MCI-C demonstrated significantly elevated betweenness centrality within the Default Mode Network (DMN), indicating that DMN hubs assumed a disproportionately important role in mediating communication across the brain. This pattern of early hub overload is consistent with the interpretation from several studies that increased functional MRI activity in preclinical and early stages of AD serves as a marker for neuronal compensation or aberrant synchronization^34, 35^. Furthermore, longitudinal functional connectivity studies show that biomarker-positive asymptomatic adults progress through a phase of increased DMN activity/connectivity before transitioning to the later network-wide decline characteristic of MCI/dementia^35, 36^. The DMN is exquisitely vulnerable to such pathological pressure due to its high metabolic demand and propensity for early amyloid and tau accumulation^15^

Our finding specifically identifies elevated betweenness centrality, a topological marker of hub overload as a key feature of this early, vulnerable network state.

Notably, the lack of a significant difference in DMN eigenvector centrality suggests a specific alteration in the DMN’s bridging function, rather than a global increase in its influence. This selective redistribution of communication load, where hubs become more essential as bridges without gaining overall network dominance, is consistent with a compensatory reorganization under early pathological pressure. This “hub overload” may, in turn, heighten the metabolic and functional vulnerability of DMN regions, potentially accelerating their decline within the AD trajectory.

### 4.4 Network Resilience: Preserved Global Robustness Despite Early Functional Reorganization

Contrary to our hypothesis, MCI-C did not differ significantly from MCI-NC in overall network resilience, as quantified by the critical drop point during targeted attack simulations. This indicates that although MCI-C exhibit functional reorganization including reduced global efficiency and increased average strength and DMN betweenness, these alterations have not yet translated into causing vulnerability of the network’s structural backbone.

Importantly, MCI-C did show a trend-level reduction in the initial largest connected component (LCC), suggesting that subtle fragility may already be present before any simulated attack. A smaller LCC at baseline implies that the functional connectome of MCI-C is slightly less globally connected, even in the absence of perturbation. Although this reduction did not meet conventional statistical significance, its consistent directionality complements the broader pattern of inefficient hyperconnectivity observed across other metrics.

Our findings of increased nodal strength but reduced global efficiency in MCI-C define a phenotype of inefficient hyperconnectivity, where stronger local coupling undermines global integration. This dissociation reflects an early compensatory phase in Alzheimer’s disease (AD) networks. De Haan et al.^37^ demonstrated through an *activity-dependent degeneration model* that networks can remain structurally robust despite declining efficiency, until the damage to these hubs triggers a “pathologic spiral” that leads to eventual network collapse. The preserved resilience observed here suggests that converters remain in an intermediate phase where high nodal activity persists, but the network has not yet reached the stage of global fragmentation predicted by the ADD (Activity Dependent Degeneration) Network resilience-focused models further clarify this window. Argiris, Stern, and Habeck^11^ network quantified resilience by probing resting-state BOLD networks under targeted attack, linking maintained integrity to cognitive reserve. A recent review by Nawani et al.^38^ reinforces this approach, proposing that resilience-based models such as targeted attack simulations and LCC decay are essential to capture the rapid, non-linear transitions in network integrity that static measures like global efficiency often obscure. They argue that early-stage networks often maintain global integration through compensatory reorganization, which masks latent fragilities and hub overload. Together, these frameworks suggest that MCI converters occupy a precarious transitional stage: they possess a functional connectome that is still globally robust yet is characterized by a “strained stability” nearing its collapse threshold.

### 4.5 Preserved Cortical Structure Reinforces the Primacy of Functional Change

Cortical thickness did not differ between baseline MCI-C and MCI-NC. This finding is consistent with the established Alzheimer’s Disease (AD) Biomarker Cascade, which posits a specific temporal sequence: molecular pathology (amyloid-beta) is followed by functional/metabolic changes (e.g., FDG-PET hypometabolism), and only later is neurodegeneration manifested as detectable cortical atrophy ^39,40^. The fact that our cohort did not exhibit statistically significant cortical thickness differences at baseline suggests that these subjects were captured at an exceptionally early preclinical stage, where the subtle, regionally specific thinning reported in other cohorts^41^ has not yet reached the threshold of statistical significance to separate converters from non-converters. This suggests our network topology findings as an earlier marker for identifying at-risk individuals.

### 4.6 Network-based prediction of MCI conversion

We evaluated multiple supervised classifiers, including logistic regression, k-nearest neighbors (KNN), support vector machines (linear and RBF kernels), and random forest, to assess the predictive utility of baseline multimodal features for conversion from high risk to mild cognitive impairment (MCI). Models were trained using repeated nested cross-validation, with hyperparameter optimization and feature selection confined to training folds. Covariate adjustment for age, sex, and years of education was applied within each training fold to prevent information leakage.

Classifier performance during outer cross-validation was moderate but consistent across models, with mean accuracies ranging from ∼57% (SVM linear) to ∼61% (logistic regression) and standard deviations around 0.12. Focusing on KNN, which showed the most balanced performance, mean outer CV accuracy was 59.6% (SD = 0.12), indicating stable predictive performance across folds. On the independent test set, accuracies ranged from 44–56%, with KNN, logistic regression, and linear SVM each achieving 55.6%, and KNN showing the highest F1 score (0.56). These modest, just-above-chance values are expected for baseline prediction, where some participants classified as non-converters may eventually progress to MCI. Additional variability may arise from inter-individual differences in cognitive reserve, lifestyle, or molecular and vascular risk factors.

Feature stability analysis was performed using KNN outer folds, quantifying the frequency with which individual features were selected across 100 repetitions. Prediction was driven predominantly by functional network topology, with global efficiency emerging as the most consistently selected feature (25.8%), followed by critical drop (19.4%), average clustering coefficient (17.8%), initial largest connected component (LCC, 16.2%), and modularity (11.2%). Together, these network measures accounted for the majority of selections, indicating that early conversion risk is primarily associated with alterations in network integration and resilience.

By contrast, p-tau217 (4.4%) and mean cortical thickness (2.8%) were selected less frequently, suggesting that molecular and structural markers provide complementary but weaker predictive information at this preclinical stage. Notably, critical drop contributed to KNN classification despite the absence of a significant group-level difference, highlighting the distinction between univariate sensitivity and multivariate predictive relevance.

Taken together, these findings indicate that subtle network-level alterations associated with conversion risk are detectable at baseline, even when overall predictive accuracy is limited. Feature selection frequency reflects risk-related network signatures rather than deterministic prediction, supporting a network-centric framework in which functional reorganization and resilience deficits precede overt structural atrophy or elevated peripheral biomarker expression.

## 5. Conclusion

This exploratory study investigated baseline functional network topology in high-risk older adults who later progressed to mild cognitive impairment (MCI). Consistent with their elevated biological risk, future converters (MCI-C) were older, had fewer years of education, and exhibited higher plasma p-tau217 levels at baseline. At the network level, MCI-C displayed a consistent pattern suggestive of “inefficient hyperconnectivity” characterized by reduced global efficiency alongside increased average nodal strength and elevated betweenness centrality within the Default Mode Network. While these topological differences did not survive strict correction for multiple comparisons, the directionally consistent effects across multiple network densities and metrics point to a meaningful, if subtle, alteration in the preclinical connectome. Supervised machine learning provided supporting evidence for the relevance of these network alterations. In a repeated nested cross-validation framework, a model based on multimodal features achieved predictive accuracy modestly above chance levels. This performance reflects the inherent difficulty of long-term prediction from a single time point in a preclinical cohort. Notably, the features most frequently selected to drive prediction were graph-theoretical metrics of integration (global efficiency) and resilience (critical drop), rather than structural or plasma biomarkers alone. This indicates that the multivariate signal of impending conversion is embedded in the pattern of functional network organization.

In summary, this study identifies a profile of demographic risk, incipient pathology, and altered functional network topology that characterizes high-risk individuals in a vulnerable preclinical state. The findings though requiring replication and validation in larger, independent cohorts generate the hypothesis that a shift towards inefficient hyperconnectivity and a specific signature of network resilience may represent an early, systems-level vulnerability that precedes overt clinical decline and detectable atrophy to understand the earliest stages of the Alzheimer’s disease cascade.

## Declarations

### Ethics approval and consent to participate

Not applicable

### Consent for publication

Not applicable

### Availability of data and materials

The datasets analyzed during the current study are available through the PREVENT-AD Open Resources via the Registered Access portal of the Canadian Open Neuroscience Platform (CONP) or the LORIS database (https://prevent-ad.loris.ca/). Access is granted to qualified researchers upon completion of a data usage agreement.

### Competing interests

The authors declare that they have no competing interests.

### Funding

The authors acknowledge funding support under strategic research projects from BITS BioCyTiH Foundation (a Section 8 not-for-profit company) hosted by BITS Pilani, supported under the National Mission of Interdisciplinary Cyber Physical Systems (NMICPS), Department of Science & Technology (DST), Government of India.

### Authors’ contributions

Hema Nawani: conceptualization, methodology, formal analysis, writing– review and editing, writing– original draft, data curation, visualization. Ranjith Jaganathan: data curation, writing– review and editing. Veeky Baths: conceptualization, project administration, resources, writing– review and editing, writing original draft, supervision, funding acquisition.

---

**Supplementary Table S1.**
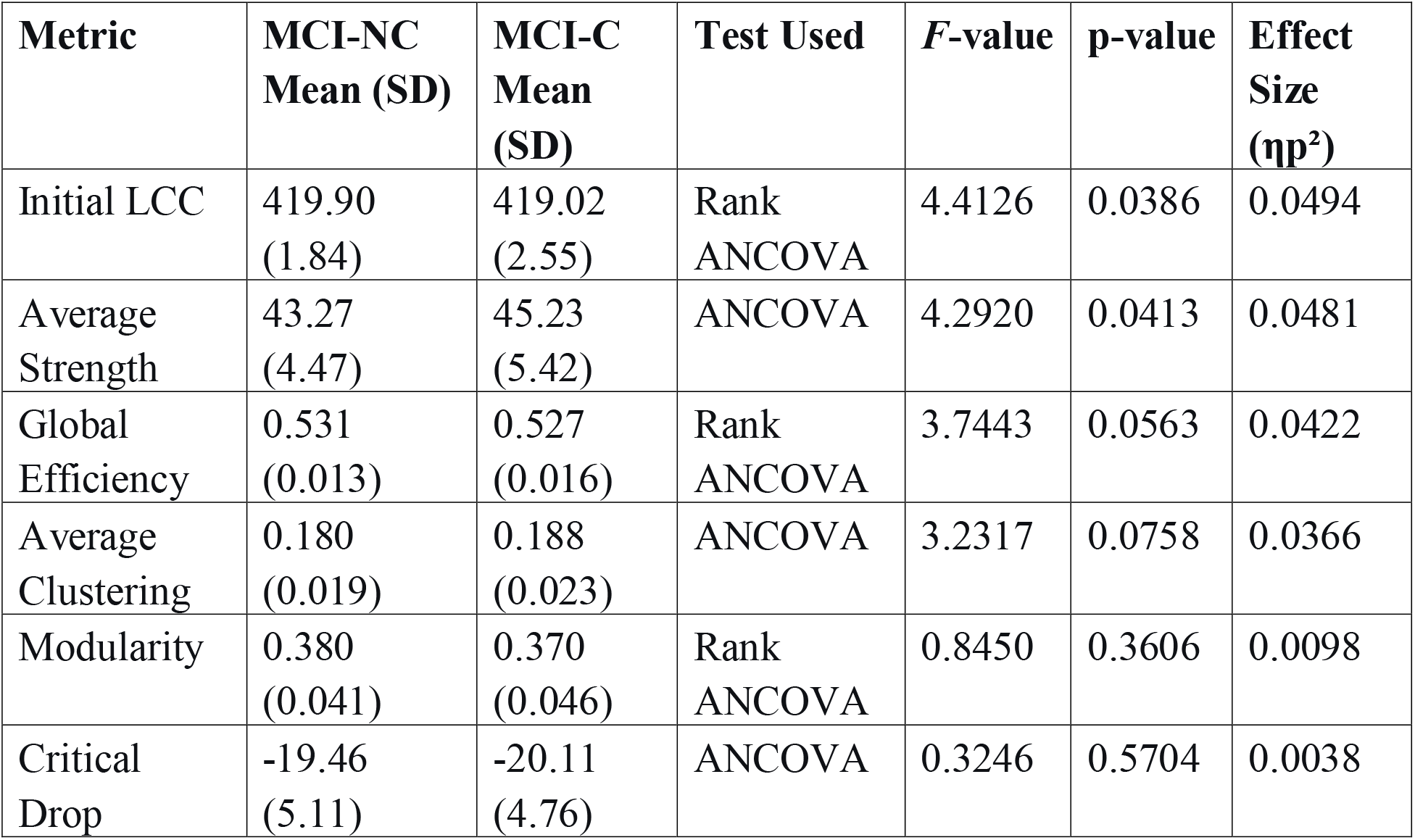
Group differences in graph-theoretical metrics at 16% network density in baseline high-risk individuals who later converted to MCI (MCI-C) and those who remained cognitively stable (MCI-NC) *ANCOVA or Rank ANCOVA controlling for age, sex, and education. Results are reported at uncorrected p < 0.05*

## Notes

### Competing Interest Statement

The authors have declared no competing interest.

