## Supplemental Table 1 for "Altered Baseline Brain Network Topology in High-Risk Individuals Progressing to Mild Cognitive Impairment"

**Supplementary Table S1. Group differences in graph-theoretical metrics at 16% network density in baseline high-risk individuals who later converted to MCI (MCI-C) and those who remained cognitively stable (MCI-NC)**

***ANCOVA or Rank ANCOVA controlling for age, sex, and education. Results are reported at uncorrected p < 0.05***

| **Metric** | **MCI-NC Mean (SD)** | **MCI-C Mean (SD)** | **Test Used** | ***F*-value** | **p-value** | **Effect Size (ηp²)** |
|---|---|---|---|---|---|---|
| Initial LCC | 419.90 (1.84) | 419.02 (2.55) | Rank ANCOVA | 4.4126 | 0.0386 | 0.0494 |
| Average Strength | 43.27 (4.47) | 45.23 (5.42) | ANCOVA | 4.2920 | 0.0413 | 0.0481 |
| Global Efficiency | 0.531 (0.013) | 0.527 (0.016) | Rank ANCOVA | 3.7443 | 0.0563 | 0.0422 |
| Average Clustering | 0.180 (0.019) | 0.188 (0.023) | ANCOVA | 3.2317 | 0.0758 | 0.0366 |
| Modularity | 0.380 (0.041) | 0.370 (0.046) | Rank ANCOVA | 0.8450 | 0.3606 | 0.0098 |
| Critical Drop | -19.46 (5.11) | -20.11 (4.76) | ANCOVA | 0.3246 | 0.5704 | 0.0038 |
